# Mechanically-mediated low-pressure cell membrane poration enables tunable intracellular delivery of traditionally impermeable cargoes in high throughput and clinical scale formats

**DOI:** 10.64898/2026.07.31.742104

**Authors:** Sophia M. Hirsch, Darby Kreienberg, Zhihui Song, Andrew Larocque, Yi Zuo, Rachel Conover, Eleni Jaecklein, Karen Gonzalez, Ain Imchen, Sean Franco, Michael A. Evans, R. Alexander Wesselhoeft, Matthew R. Swiatnicki, Hicham Zegzouti, Kirill Chesnov, Marius Wernig, Neil S. Dhawan, Cameron Pye, Scott M. Loughhead, Jacquelyn L. Sikora Hanson, Alec Barclay, Armon Sharei

## Abstract

Traditional intracellular delivery methods suffer from cargo inefficiencies, non-linear delivery kinetics, and cellular trauma. We designed a mechanically-mediated poration platform governed by deterministic, passive diffusion operating at low pressure which enables dose-dependent, cargo-agnostic intracellular delivery and preserves cellular homeostasis; key advantages include high-fidelity multiplexing, transient cell engineering, and direct-to-biology live-cell target engagement enabling development of novel intracellular delivery applications across drug discovery and cell therapy. We demonstrate examples including live-cell DEL discovery and MOA studies, complex and rapid cell therapy manufacturing, and assay development in sensitive primary cell types.

## Main

The precise delivery of exogenous macromolecules into primary cells remains a fundamental bottleneck in advanced cell therapies and early-stage drug discovery^1–3^. While viral vectors and lipid-based transfection systems are widely utilized, they are inherently constrained by limited cargo capacity, immunogenicity, and lineage-specific compatibility^1,4^. Physical delivery methods offer a broader cargo range but are difficult to scale, lack independently tunable operational parameters, and frequently induce severe cellular stress^1–6^. Mechanical methods, such as microinjection or microfluidic mechanoporation, address cell health concerns but have throughput limitations and scaling challenges^1,3,7^. In drug discovery, the plasma membrane permeability barrier frequently necessitates the use of surrogate models or extensive chemical modification of lead candidates, obscuring true biological activity and delaying therapeutic development^8,9^.

To overcome these limitations, we engineered a mechanically-mediated poration platform designed for low-pressure, high-throughput intracellular delivery (Fig 1a). The system utilizes ultra-thin (100 nm to 2 μm) high-porosity silicon membranes featuring defined arrays of uniform circular micro-pores integrated into a rigid silicon-on-insulator (SOI) support framework (Fig. 1b). This de novo architecture provides the structural stability required for high-flow applications while maintaining a streamlined fluidic pathway that eliminates stagnation zones and minimizes hydraulic resistance, reducing challenges associated with microfluidic delivery methods such as high pressure shear and clogging^2,3,7^. The platform operates at low pressures (typically ≤10 psi), allowing for seamless integration with standard pump-based systems and eliminating external gas canister requirements. The 2D design enables effective scaling from research to higher cell throughput clinical or screening applications. Pore diameters ranging from 3 – 15 μm accommodate diverse cell diameters and membrane elasticities. Here, we demonstrate the utility of this mechanically-mediated poration platform (termed “boosting”) to provide a robust framework for rapid, perturbation-free intracellular delivery of diverse and traditionally “impermeable” cargoes proportional to concentration across automated, high-throughput drug discovery workflows and scalable, closed-system clinical cell manufacturing pipelines.

**Figure 1.**
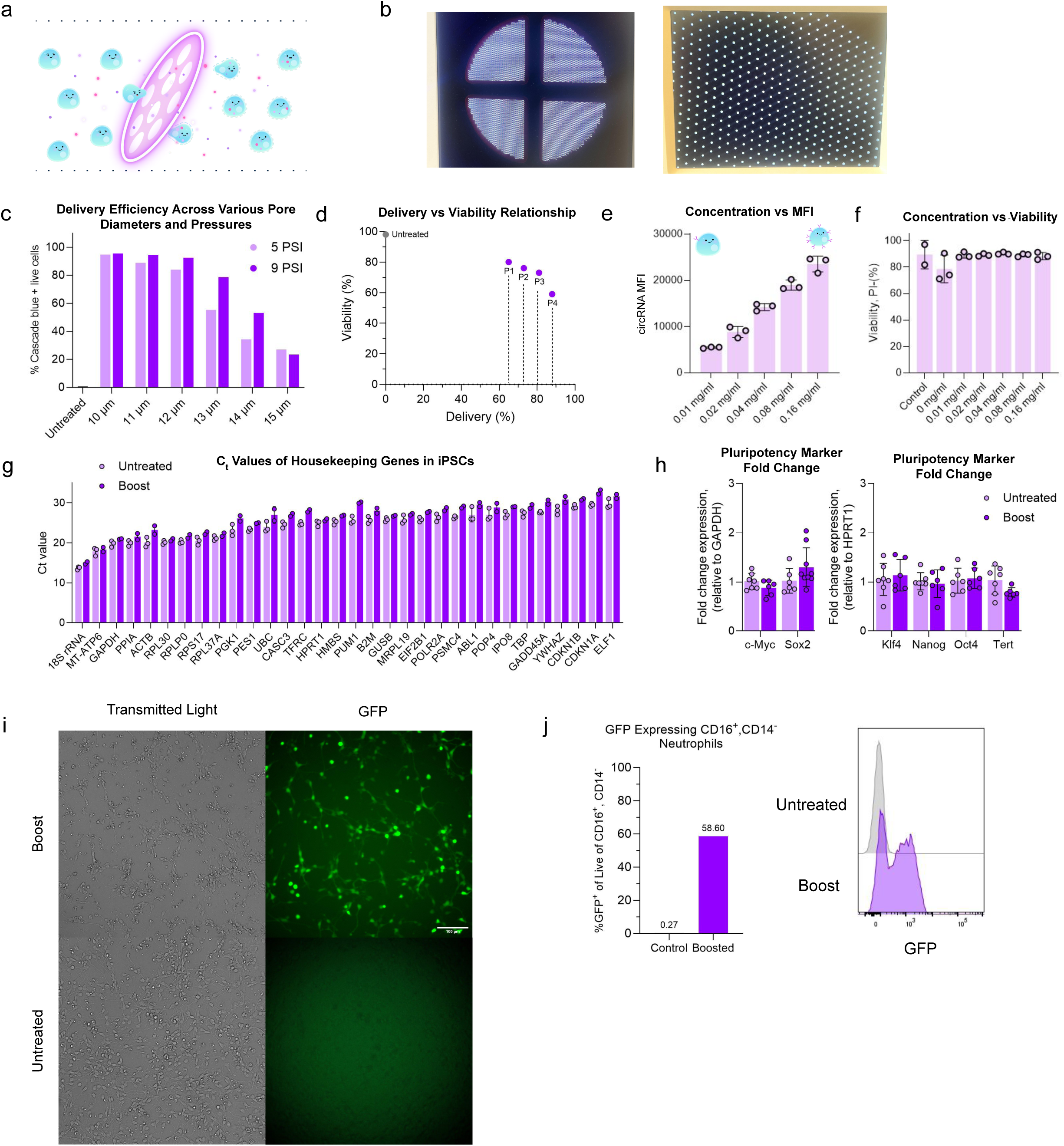
Characterization of mechanically-mediated poration and cellular health for use in sensitive and intractable cell types. **a**, Mechanism of mechanically-mediated poration (“boost”), where cells are driven by applied pressure through pores smaller than the cell diameter to achieve uniform, diffusive cytosolic uptake. **b**, Pictures of the silicon-based poration framework featuring an ultra-thin membrane (100 nm to 2 μm) with uniform circular pores supported by a rigid scaffolding structure. **c**, Delivery efficiency (percentage of cargo-positive HeLa cells) as an empirical function of applied pressure and pore diameter; higher efficiency is typically modulated by increasing operational pressure or selecting a reduced pore size. **d**, Delivery efficiency and post-poration viability are inversely correlated as demonstrated in iPSCs by higher 3 kDa dextran delivery and lower viability as operational pressure is increased. **e**,**f**, Dose-dependent titration of intracellular delivery (mean fluorescence intensity, MFI) scaling with initial cargo concentration (**e**) without impacting post-delivery viability (**f**) (shown for mbIL-2 circRNA delivery to HeLa cells). **g**,**h**, RT-qPCR quantification of housekeeping genes (**g**) and pluripotency markers (**h**) in iPSCs 48 hours post-poration, demonstrating transcriptional profiles indistinguishable from controls (n = 6 from two independent experiments). **i**, High efficiency (∼75%) and diffuse *GFP* mRNA expression in neural progenitor cells (NPCs) following mechanical delivery as compared to controls which were not boosted. Scale bar, 100 μm. **j**, GFP expression in primary neutrophils following mRNA delivery, comparing untreated control cells against platform-porated (“boost”) conditions.

Unlike conventional intracellular delivery methods that rely on unpredictable or binary uptake mechanisms^1,10^, mechanically-mediated poration ensures immediate, uniform cargo entry driven entirely by passive cytosolic diffusion^10^. This influx is driven by the interplay between cell speed (applied pressure) and pore size relative to cell diameter (Fig. 1c). Consequently, cargo delivery efficiency operates as a precise, dose-dependent response inversely correlated with viability, where higher intracellular delivery efficiencies are achieved by elevating operational pressure or reducing pore size with predictable viability costs (Fig. 1d). Once baseline fluidic parameters are established, the precise intracellular dose can be further titrated by adjusting the initial cargo concentration (Fig 1e-f, Extended Data Fig. 1a-d). Benchmarking across immortalized lines, induced pluripotent stem cells (iPSCs), and primary human peripheral blood mononuclear cell (PBMC) subtypes yielded consistent high-efficiency uptake and robust survival across diverse lineages following simple optimization of the empirically tunable parameters (Extended Data Fig. 1e-p). Our approach contrasts sharply with the stochastic or binary delivery profiles characteristic of physical and biological delivery modalities which offer fragmented, non-linear parameter control and are plagued by endosomal entrapment challenges^1,10,11^, establishing this platform as a reliable framework for predictable and precise delivery of diverse cargoes across a wide array of sensitive cell types (Extended Data Fig. 1q).

Mechanoporation technologies achieve robust intracellular delivery while uniquely preserving cellular homeostasis, effectively eliminating cellular trauma associated with high-voltage electroporation^6,7^. Cellular preservation following processing on our platform (termed “boost”) was evaluated via RT-qPCR transcriptomic analysis of housekeeping genes and pluripotency markers in iPSCs, revealing expression profiles indistinguishable from untreated controls (Fig. 1g-h). These data confirm maintenance of the cellular native state and lineage fidelity which is essential for downstream therapeutic applications or maintaining proper target scaffolding in drug discovery screens. To demonstrate this advantage, we delivered mRNA and circular RNA (circRNA) with robust expression in traditionally “undeliverable” neuron progenitor cells (NPCs) and primary neutrophils (Fig. 1i-j), which are lineages notoriously recalcitrant to electroporation due to high sensitivity to ionic flux and electrical stress^1,12^. NPCs exhibited normal morphology (Fig. 1i), while neutrophils retained cytokine-mediated migration response (Supplemental videos 1,2). By maintaining native intracellular structural and functional integrity, our platform expands cell engineering opportunities in highly sensitive and underutilized cell populations.

A defining advantage of the diffusive uptake characteristic of boosting is the ability to achieve simultaneous, high-fidelity co-delivery of multiple molecular species. In traditional multi-modal cell engineering with stochastic delivery systems, a central challenge is compounding inefficiency and yield loss^13,14^, which drop precipitously with increasing number of target modalities, as success probability is the product of individual cargo efficiencies. Our platform bypasses this statistical limitation by enabling simultaneous, diffusive cytosolic uptake through transiently formed pores, where intracellular cargo volume is proportional to initial concentration and molecular size while preserving the stoichiometry of the starting cargo mixture within the intracellular environment, resulting in multiplexed delivery that tracks with the high baseline efficiency of the platform rather than the independent probability of multiple entry events (Extended Data Fig. 2a-g). A single-step workflow enabled robust engineering of multiple modalities to achieve simultaneous CRISPR-mediated gene editing, high-magnitude mRNA expression, and intracellular tagging (Fig. 2a-c). This multiplexing proficiency provides a powerful framework for complex biological modeling while completely avoiding cargo-specific biases, staggered expression timelines, metabolic burden, and reduced multi-modal yield associated with sequential transfection protocols.

**Figure 2.**
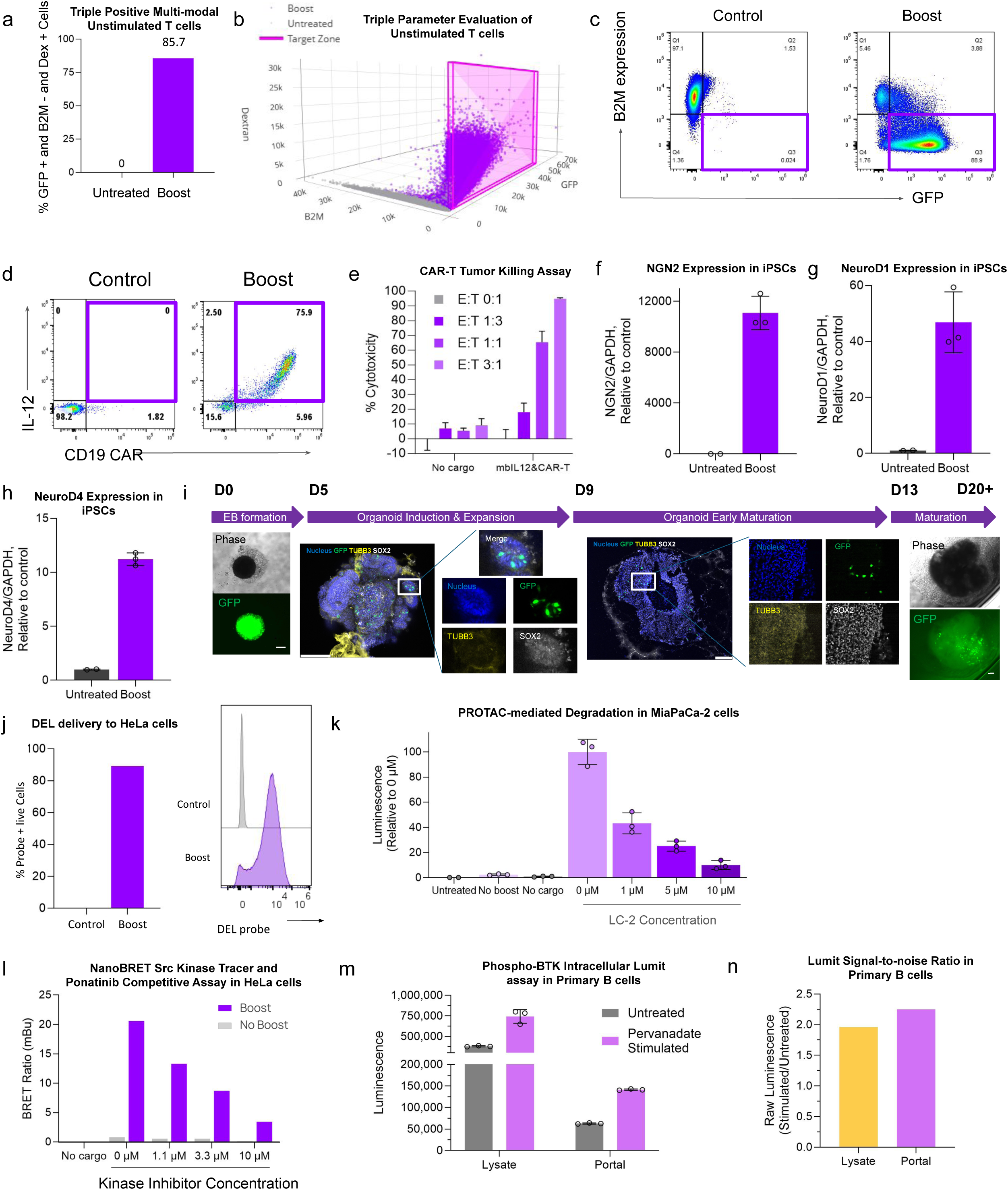
Unlocking novel applications of mechanically-mediated, low-pressure poration: efficient multi-modal and specialized cell engineering and delivery of traditionally impermeable molecules. **a-c**, Simultaneous co-delivery of *GFP* mRNA, CRISPR RNPs targeting *B2M*, and 3 kDa dextran in primary human T cells, illustrating highly efficient, single-step multiplexed cell engineering (simultaneous protein expression, genomic editing, and cell labeling). Shown are representative quantification of triple positive cells (**a**), 3D overlay plot of surface B2M, GFP expression, and intracellular dextran in boosted vs untreated with target area defined (B2M-/GFP+/dextran+) (**b**), and 2D scatter plot depicting surface B2M and GFP expression. **d,e**, Multi-modal expression kinetics in primary human T cells transiently engineered with circRNA encoding a CD19-targeted chimeric antigen receptor (CAR) and membrane-bound interleukin-12 (mbIL-12) (**d**), and corresponding downstream cell-mediated tumor killing dynamics (**e**). **f**-**h**, Directed differentiation of human iPSCs into early neurons following *Ngn2* mRNA boost. RT-qPCR quantification demonstrates upregulation of early neuronal markers *Ngn2* (**f**), *NeuroD1* (**g**), and *NeuroD4* (**h**) relative to *GAPDH* compared to untreated controls. **i**, Maturation of cerebral organoids demonstrating sustained GFP expression following *GFP* saRNA boost to human iPSCs using a standard 20-day differentiation protocol. Scale bar, 200 μm. **j**, Intracellular delivery of DNA-encoded libraries (DELs) to HeLa cells. DEL concentration following platform-mediated delivery to PBMCs correlates linearly with diluted DEL input as quantified by sequencing (Spearman R^2^=0.93). **k**, Live-cell kinetic monitoring of PROTAC-mediated degradation. Mechanical-mediated delivery of LgBiT protein into MiaPaCa-2 cells (endogenously tagged with HiBiT at the *KRAS(G12C)* locus) yields a dose-dependent decrease in luminescent signal corresponding to degradation by the PROTAC LC-2. **l**, Dose-dependent BRET signal decrease indicating competitive displacement by the Src kinase inhibitor Ponatinib in NanoLuc-Src-expressing HeLa cells following intracellular loading of a traditionally impermeable kinase tracer via the platform. **m**,**n**, Live-cell intracellular phospho-BTK detection in activated human B cells. Mechanical delivery of four phospho-BTK antibodies following pervanadate stimulation yields a robust intracellular phosphorylation signal (**m**) with a signal-to-noise ratio comparable to traditional lysate-based detection (**n**).

Direct cytosolic access for multiple modalities paired with native state preservation is particularly transformative for advancing cell therapies by enabling complex cell engineering within a homeostatic environment; use of transient cargoes, such as circRNA^15–17^ can drive profound biological responses without the genomic integration or sustained cellular stress characteristic of viral and DNA-based methods. We leveraged this capability to transiently engineer primary human T cells with circRNA constructs encoding a CD19-targeted chimeric antigen receptor (CAR) and membrane-bound interleukin-12 (mbIL-12). This single-step multi-modal delivery yielded rapid onset and high-magnitude co-expression of both therapeutic modalities (Fig. 2d), leading to enhanced tumor cell killing efficacy (Fig. 2e) while circumventing permanent genomic manipulation and long-term cellular perturbation. We extended this transient modulation paradigm to neurogenesis, demonstrating that mRNA delivery to iPSCs can drive early neurodevelopment (Fig. 2f-h). Further, boosting GFP self-amplifying RNA (saRNA) results in long-lasting GFP positive cells and does not disrupt lineage commitment or structural scaffolding during a 20-day cerebral organoid maturation protocol (Fig. 2i). By establishing a highly efficient, non-viral methodology for immediate macromolecular expression, this transient approach provides a platform for robust, clinically relevant methodology to accelerate functional oncology modeling, target validation in complex lineages, and the accelerated manufacturing of next-generation cellular therapeutics.

The ability to bypass the plasma membrane permeability barrier and inefficiencies of endosomal escape opens new opportunities within early-phase drug discovery by providing a universal pathway for cytosolic delivery of diverse molecular libraries and analytical tools. For early screening, boosting enables cytosolic entry of traditionally “impermeable” modalities, including small molecules^18^, peptides (Extended Data Fig 1a-d), and DNA-encoded libraries (DELs) (Fig. 2j & Extended Data Fig 2h); this provides potential to both accelerate timelines and transform drug candidate identification and characterization in the context of native intracellular scaffolding, even within primary cells rather than engineered lines. While traditional mechanism-of-action (MOA) studies depend on oversimplified cell-free lysates, harsh chemical detergents or surrogate immortalized cell lines, mechanically-mediated poration enables direct target evaluation within a cell’s native scaffolding. We validated three distinct implementations of this approach, serving as a foundation for a myriad of designs that capitalize on live-cell native MOA study: first, delivering the LgBiT protein directly into cells endogenously tagged with HiBiT facilitated precise, real-time kinetic monitoring of PROTAC-mediated target protein degradation (Fig. 2k); second, non-viral intracellular loading of an otherwise impermeable Src kinase tracer resulted in dose-dependent competitive displacement by Src small-molecule inhibitors (Fig. 2l, Extended Data Fig. 2i,j); and lastly, intracellular co-delivery of the four antibody components of the Lumit immunoassay^19^, consisting of a pair of primary antibodies against total BTK and phospho-BTK (Y223), together with Lumit secondary antibodies (anti-mouse-LgBiT and anti-rabbit-SmBiT), enabled live-cell, wash-free detection of intracellular kinase phosphorylation following pervanadate stimulation with signal-to-background ratios comparable to conventional lysate-based Lumit workflow^19^ (Fig. 2m,n & Extended Data Fig. 2k,l). By eliminating artificial proxy models and preserving native target conformations, post-translational modifications, and cellular pathways, this direct-to-biology framework compresses validation timelines from months to hours while delivering deeper, physiologically accurate insights into candidate drug mechanisms.

The porous 2D architecture of this platform enables full translational utility across operational scales with simple scaling of design footprint to accommodate throughput requirements, providing a unified framework that bridges the gap between early-stage discovery and large-scale therapeutic manufacturing. For high-throughput (HT) discovery applications, the low-pressure requirements of the silicon membrane enable seamless integration with existing automated cell dispensing instrumentation (Fig. 3a & Extended Data Fig. 3a). This approach leverages a distinct attribute of mechanically-mediated poration: membrane closure can take up to a minute or more^1,10^, allowing for diffusion of cargo into the cell after the poration event, and enabling a workflow in which cells can be mechanoporated during dispensing into a plate which is pre-loaded with cargo (Fig. 3b, Extended Data Fig. 3b-e). This principle provides the foundation for automated delivery of cargo in 96- and 384-well formats, achieving uniform cargo delivery across thousands of independent samples without the need for specialized high-pressure infrastructure (Fig. 3c-f). This ease of integration and scaling allows the platform to be deployed within standard liquid-handling workflows with minimal effort, providing the foundation for novel assay applications including CRISPR screens, traditionally impermeable molecule screens, use of primary cells in screening assays, and multi-modal screening capabilities.

**Figure 3.**
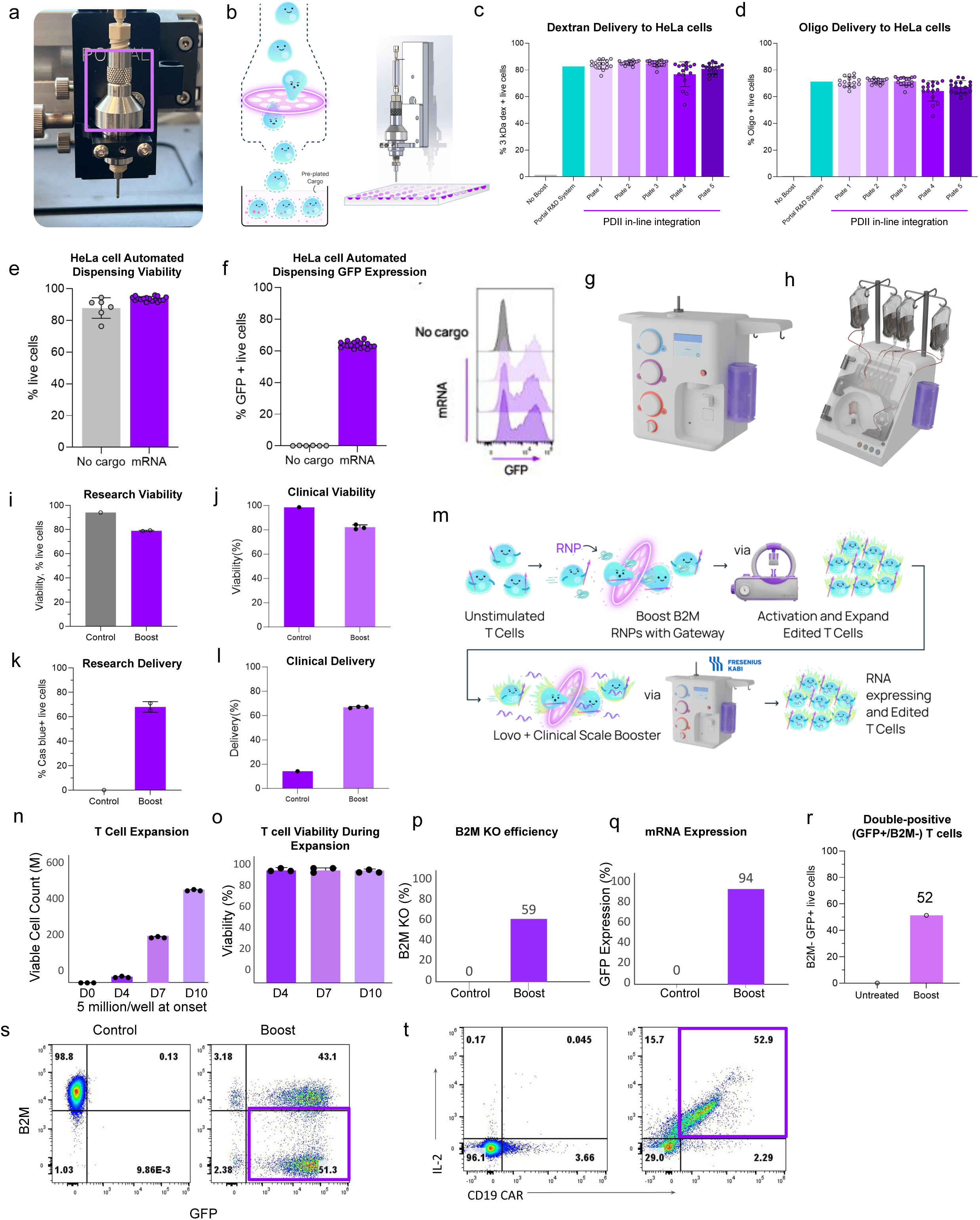
Scalability of the mechanically-mediated platform for high-throughput discovery and clinical manufacturing. **a**,**b**, Integration of the mechanically-mediated poration platform with standard automated liquid handling systems, showing a representative hardware configuration with the HighRes PreciseDrop II (PDII) (**a**) and a schematic depicting cell transit through the porous membrane for direct dispensing into pre-plated cargo (**b**). **c**,**d**, Performance consistency across multiple plates using automated fluidic dispensing, evaluated by introducing 3 kDa dextran (**c**) or oligonucleotides (**d**) into HeLa cells via the cargo pre-plating method using the PDII system. **e**,**f**, Cell viability (**e**) and delivery efficiency (**f**) following integrated mechanically-mediated poration and automated dispensing of HeLa cells into pre-plated *GFP* mRNA. **g**,**h**, Schematic of the closed-system clinical scaled-up architecture utilizing sterile-weld integration with Fresenius Kabi LOVO cell processing instrumentation (**g**) and Thermo Fisher Scientific CTS Rotea Counterflow Centrifugation System (**h**). **i**-**l**, Comparison of cell viability (**i**,**j**) and delivery efficiency (**k**,**l**) across micro-scale (research scale, 5 x 10^6^ cells, **i**,**k**) and milli-scale (clinical scale, 5 x 10^8^ cells, **j,l**) processing volumes (n = 2 independent biological replicates for research scale, n = 3 for clinical scale). **m**-**s**, Functional validation of human primary T cells processed in a multi-day, multi-scale clinical manufacturing workflow (**m**) incorporating sequential research-scale RNP editing and clinical-scale mRNA delivery. Data represent mean ± s.d. from three independent biological replicates. **n**,**o**, Longitudinal cell expansion (**n**) and post-poration viability (**o**) over a 10-day activation period following initial benchtop-scale delivery of CRISPR RNPs to unstimulated T cells. **p**-**s**, Quantitative multi-modal engineering efficiencies at study completion following *GFP* mRNA delivery to activated T cells within the closed, integrated clinical-scale LOVO system. Data show *B2M* knockout (KO) efficiency (**p**), GFP expression (**q**), and resultant dual-engineering (*B2M* KO and GFP expression) efficiency (**r**) with scatter flow plot comparing B2M-/GFP expressing cells between control and boost groups (**s**). **t**, Representative flow cytometry plots demonstrating multi-modal engineering of human T cells directly in whole blood via research-scale delivery of multiplexed circRNA for a CD19 CAR construct and mbIL-2 via the research-scale platform.

For therapeutic applications requiring substantial cell volumes, the platform architecture scales linearly across orders of magnitude to accommodate processing billions of cells through a closed-system configuration designed for sterile-weld integration into clinical-grade processing instrumentation, including the Fresenius Kabi LOVO (Fig. 3g) and CUE systems and Thermo’s Rotea system (Fig. 3h). This integrated workflow enables the rapid production of engineered cell therapies, such as CAR-T cells, with efficiencies and viabilities that mirror those observed at research scale (Fig 3i-l, Extended Data Fig. 3f-j). To evaluate the platform’s compatibility with a multi-step clinical manufacturing process, we performed a longitudinal study involving the sequential engineering of primary human T cells over a 10-day expansion period (Fig. 3m). The workflow commenced with research-scale delivery of CRISPR RNPs for B2M editing in unstimulated T cells, followed by a standard clinical activation protocol, with robust, >80-fold expansion (Fig. 3n) and high viability up to clinical-scale densities (Fig 3o). The population then underwent a second, clinical-scale boost of GFP mRNA using a porous membrane integrated in-line with the LOVO system, resulting in high-efficiency editing (∼60% B2M knockout, Fig. 3p, Extended Dat Fig. 3k) and near-universal mRNA expression (94% GFP+, Fig. 3q) with double-positive cell rate matching the lowest efficiency (52% B2M-/GFP+, Fig. 3r,s) at study completion. A separate study demonstrated proof of concept for engineering human T cells in whole blood with circRNA for CD19 and IL-2 for a more rapid protocol that eliminates the need for T cell isolation (Fig. 3t). The sustained performance of T cells across multiple deliveries and orders of magnitude in cell volume along with ability to manufacture cells within whole blood demonstrates the platform’s unique capacity to serve as a unified “plug-and-play” solution for various complex, longitudinal therapeutic manufacturing.

In summary, this mechanically-mediated poration platform introduces a deterministic, low-pressure architecture that replaces traditional stochastic vectors to prioritize preservation of the cellular native state while seamlessly integrating into automated and clinical instrumentation. By decoupling intracellular delivery from conventional biological and physical constraints, this fluidic interface unlocks unprecedented engineering capabilities, including: multi-modal engineering of highly sensitive, historically intractable lineages; direct loading of traditionally impermeable molecules; and, linear operational scaling across multiple orders of magnitude. Ultimately, by bringing traditionally restricted assay tools and molecular modalities directly into unperturbed cellular environments, this plug-and-play framework dramatically accelerates both early-phase live-cell drug discovery and validation, as well as large-scale cell therapy manufacturing.

## Methods

### Ultra-thin silicon membrane design and fabrication

Micro-pore arrays were engineered and fabricated from SOI wafers, with the geometric configuration optimized to balance structural integrity with high volumetric processing flow rates under positive pressure. The initial substrates comprised a thin silicon filtering layer (1 – 50 μm), a buried silicon oxide stop layer (0.1 – 10 μm), and a thicker silicon support layer (100 – 500 μm). To define the mechanically-mediated poration apertures, the active filtering layer was photolithographically patterned and etched linearly via deep reactive ion etching (DRIE). Etching terminated precisely at the sacrificial buried silicon oxide stop layer to guarantee highly uniform pore geometry and tolerance across the array (circular pore diameters ranging from 3 to 15 μm). Following pore formation, the substrate was stripped of photoresist and cleaned.

To prevent mechanical failure of the ultra-thin membrane during operation, a secondary support structure template was patterned onto the opposing support layer. DRIE was utilized to etch through the support layer up to the buried oxide interface, creating an integrated framework of supporting structure, designed to mask a minimal portion of the total porous area to optimize volumetric processing flow rates. The exposed sections of the sacrificial oxide layer were selectively removed via reactive ion etching to establish open, continuous fluidic channels through the micro-pore arrays. Finished silicon wafers were diced into standardized operational formats (ranging from 2 x 2 mm^2^ to 14 x 14 mm^2^ chips) scaled for desired throughput and matched to downstream cartridge housings.

### Silicon membrane integration into automated dispensing instrumentation

To enable high-throughput mechanically-mediated poration workflows, silicon membranes were integrated in-line using custom housings at the output of the fluidic pathway of commercially available, automated dispensing instruments including the Certus Flex (Nnano/Gyger), Precise Drop II^TM^ (PDII) (HighRes), WellJet (Integra), and GNF Systems: One Tip Dispenser. During standard operation of the instruments, cells passed through the installed membrane, underwent mechanically-mediated poration, and were subsequently dispensed into a 96- or 384-well plate. Cargo was pre-plated into the 96- or 384-well collection plate for post-poration diffusive uptake. For constant-pressure-driven instruments such as the Certus Flex and PDII, onboard pneumatic pressure drove cell transit through the micro-pores. Solenoid valves on the automated dispensing instruments individually dispensed porated cells into separate wells. For peristaltic-based integrations, including the Integra WellJet, instrument roller speed was adjusted to optimize cell transit velocity and mechanically-mediated poration performance.

### Silicon membrane integration into clinical cell processing instrumentation

To adapt the platform for clinical-scale applications, silicon membranes were integrated in-line with the fluidic pathways of clinical cell processing instrumentation, including the LOVO Cell Processing System (Fresenius Kabi) and the CTS Rotea Counterflow Centrifugation System (Thermo Fisher Scientific). This integration utilized the instruments’ onboard peristaltic pumps to drive cells through the integrated silicon membranes. Custom housings designed for this integrated mechanically-mediated poration were sterile welded into disposable cell processing kits and connected to instrumentation. Variable roller pump speeds were adjusted to optimize cellular transit velocity and poration efficiency. Following processing, porated cells were collected directly into sterile-connected cell processing bags within the closed fluidic system.

Silicon membrane housing assembly was designed to accommodate high-volume cell suspensions. To ensure uniform transit velocity, housings incorporated upstream pressure-dampening components which minimized flow oscillations induced by peristaltic pump rollers. The housing assembly also included an upstream pre-filtration module to capture cell aggregates, thereby preventing micro-pore clogging and extending membrane operational lifespan.

### Mechanically-mediated poration

Intracellular cargo delivery was achieved by passing cell suspensions through porous silicon membranes utilizing various micro-pore architectures designed for scale and integration type and driven by positive pneumatic pressure. Target cells were harvested from culture or isolated from primary source, washed, and resuspended to generate a single-cell suspension in minimal cell culture medium or a lineage-specific buffer. Final cell densities were optimized by lineage, ranging from 1 x 10^6^ to 5 x 10^7^ cells/mL.

For pre-mix delivery workflows, cargo(es), including fluorescently tagged dextran polymers, mRNA, circRNA, siRNA, saRNA, macrocycles, proteins, or DEL probes, were added directly to the cell suspension prior to processing. CRISPR RNPs were pre-complexed by mixing Cas9 protein with guide RNA (gRNA) and incubating at room temperature (RT) for at least 15 minutes prior to cell introduction. For HT post-mixing applications, cargo was pre-distributed directly into a multiwell collection plate.

The mechanically-mediated poration process was governed by micro-pore size and processing transit velocity, driven by applied pressure, which were carefully selected and optimized for each cell lineage. Silicon membrane pore diameters (ranging from 3 to 15 μm) were selected based on target cell diameter. For each lineage, optimal operational windows were determined empirically by using a parameter matrix crossing multiple pore sizes with a range of applied pressures.

Processing transit velocity was tuned using positive pressure across different pump and hardware interfaces scaled to the workflow application. At research scale, the cell and cargo suspension was loaded into a cartridge containing the porous silicon membrane; a pneumatic pump and regulator were then used to drive cell transit at a constant applied pressure maintained in a range from 2 to 12 psi. For HT workflows integrated into a dispensing instrument, the cargo-absent cell suspension was loaded into the designated instrument reservoir. The instrument pump system drove the cells through a membrane-containing aperture coupled to the dispensing arm, and porated cells were collected in the cargo-containing multiwell plate. Clinical, instrument-integrated workflows utilized a large-format cartridge which was sterile-welded in-line with tubing on standard cell processing equipment. Pre-mixed cells and cargo were introduced via bag or syringe interface, and driven through the cartridge using the instrument’s internal pump assembly.

Following micro-pore transit, the processed cell suspensions were collected and incubated statically at RT for at least 30 seconds to permit passive cytosolic diffusion of the cargo before membrane resealing. The delivery environment was subsequently diluted with pre-warmed complete growth medium appropriate for the lineage, and cells were transferred to downstream functional assays or standard culture vessels.

All research-scale procedures and all cell or cargo preparations were performed under sterile conditions within a laminar flow hood. Any preparations transferred to instrumentation outside of the hood were maintained in closed containers to ensure sterility.

### Microscopy imaging

Following overnight culture post-GFP mRNA delivery via mechanically-mediated poration, live neurons were imaged directly in their culture vessels using a Leica DMI8 inverted microscope. To assess cellular morphology and mRNA expression, paired phase contrast and green fluorescence images were acquired across representative fields of view. All imaging was performed using standard GFP filtration and captured via Leica Application Suite X (LAS X) software.

### Flow cytometry analysis

Cells were harvested, washed with phosphate-buffered saline (PBS), and resuspended in flow cytometry staining buffer (PBS supplemented with 2% fetal bovine serum (FBS) and 2 mM ethylenediaminetetraacetic acid (EDTA)). Where applicable, cells were incubated with fluorophore-conjugated antibodies for 20 to 30 minutes at RT in the dark, in accordance with the manufacturer’s recommendations. Following staining, cells were washed twice with staining buffer and resuspended for data acquisition. Viability was assessed using a Fixable Live/Dead viability dye (Thermo) or Propidium Iodide (PI). Where applicable, compensation matrices were established using single-stained compensation controls to ensure accurate gate placement. Flow cytometric acquisition was performed utilizing an Attune NxT Flow Cytometer (Thermo Fisher Scientific).

High-dimensional data analysis was performed utilizing FlowJo software (BD Biosciences). Hierarchical gating strategies were established using unstained or untreated controls as appropriate, and applied consistently across all experimental samples. Analytical gates defining successful intracellular delivery were established using baseline controls in which cells were mixed with the target cargo but not porated. Results are presented as the percentage of positive cells and/or median fluorescence intensity (MFI), as specified.

### RT-qPCR gene expression analysis in iPSCs

Human iPSCs (SCTi003-A; STEMCELL Technologies) were maintained in eTeSR (STEMCELL Technologies) supplemented with 10% CloneR2 (STEMCELL Technologies) and 1% Penicilin/Streptomycin (Corning) on a Matrigel substrate (Corning) at 37 °C under 5% CO_2_. Cells were lifted into single cell suspension using Accutase (Thermo), passed through a 40 μm cell strainer (VWR), and resuspended at a final density of 2 x 10^7^ cells/mL in basal eTeSR media. Cells were boosted for the intracellular delivery of 3 kDa Cascade blue dextran (Thermo) and *GFP* mRNA (TriLink). Following micro-pore transit, the engineered cells were seeded on matrigel substrate and cultured as described above. Successful cargo delivery was confirmed via flow cytometry (>70% of live cells).

Following a 48 hour incubation period, total RNA was extracted using the RNeasy kit (Qiagen) in accordance with the manufacturer’s instructions. RNA concentration and purity were determined by spectrophotometric analysis prior to reverse transcription. Complementary DNA (cDNA) was synthesized from purified RNA utilizing the SuperScript™ IV First-Strand Synthesis System (Thermo Fisher Scientific) in accordance with the manufacturer’s protocol.

Quantitative real-time PCR (RT-qPCR) was performed using TaqMan™ Gene Expression Assays (Thermo Fisher Scientific) on a QuantStudio™ 7 Real-Time PCR System (Thermo Fisher Scientific). Housekeeping genes were assayed using TaqMan Array Human Endogenous Control plates (Thermo Fisher Scientific). Reactions were prepared following the manufacturer’s recommendations and run under standard thermal cycling conditions with three technical replicates per sample. Relative gene expression levels were quantified using the comparative C_t_ (2^−ΔΔCt^) method, with target expression normalized to the endogenous reference genes HPRT1 or GAPDH, and relative sample expression values calculated by comparison to the designated experimental controls.

### Neutrophil migration assay

Primary human neutrophils in whole blood from healthy donors (NaHep anticoagulant) were maintained in whole blood for mechanically-mediated loading of *GFP* circRNA and dextran through recovery steps. Following post-poration recovery, cells were cultured for 3 hours at 37 °C under 5% CO_2_ in X-VIVO 15 medium (Lonza) supplemented with 5% human AB serum (MilliporeSigma), 2 mM L-glutamine, 1X NEAA, and 55 μM 2-mercaptoethanol. Post-incubation, cells were collected and pelleted (300 x g for 5 min). To remove contaminating erythrocytes, red blood cells (RBCs) were lysed via two consecutive 5 minute incubations in 1 mL ACK lysis buffer (Gibco) at RT with gentle shaking.

For functional migration assessment, cells were seeded onto fibronectin-coated 96-well plates and allowed to settle for 10 minutes prior to imaging. Chemotaxis was induced by addition of chemoattractant formyl-methionyl-leucyl-phenylalanine (fMLP) to the media. Resulting migration of GFP^+^ cells was tracked via time-lapse microscopy with images captured every 5 seconds using an inverted Nikon Ti2-E microscope (Nikon Instruments).

### Organoid development

Human iPSCs (SCTi003-A; STEMCELL Technologies) were used to generate cerebral organoids following the STEMdiff™ Cerebral Organoid Kit protocol (STEMCELL Technologies) with minor modifications. Following mechanically-mediated poration for the delivery of GFP saRNA, cells were resuspended in Organoid Formation Medium supplemented with 10 μM Y-27632 (ROCK inhibitor) at a final density of 6 x 10^4^ cells/mL. Suspensions were seeded at 150 μL per well into U-bottom 96-well plates pre-treated with Anti-Adherence Rinsing Solution to prevent cell adhesion and promote embryoid body formation and maintained at 37 °C under 5% CO_2_. On Day 3, each well received an additional 150 μL of Organoid Formation Medium. On Day 5, the culture medium was fully replaced with 150 μL of Induction Medium, and organoids were incubated statically for an additional 2 days. On Day 7, organoids were embedded in growth factor-reduced Matrigel and transferred to Expansion Medium for 3 days to support neuroepithelial budding and expansion. On Day 10, the culture environment was transitioned to Maturation Medium, which was subsequently refreshed every 3 to 4 days for the remaining duration of the culture.

### Tumor killing assay in T cells

Human T cells were isolated from PBMCs using the EasySep Human T Cell Isolation Kit (STEMCELL Technologies) following PBMC isolation from fresh peripheral whole blood from healthy donors using a standard Ficoll-based separation protocol^20^. T cells were activated using CD3/CD28 Dynabeads (Thermo), then engineered for the transient expression of a CD19-targeting CAR and mbIL-12 by boosting cell suspensions with pre-mixed separate circRNAs. Post-poration, engineered cells were rested for 2 hours at 37 °C under 5% CO_2_ in X-VIVO 15 medium with 5% human AB serum (MilliporeSigma), 2 mM L-glutamine, 55 μM 2-mercaptoethanol, 1X NEAA, 200 U/mL IL-2, 10 ng/mL IL-7 (PeproTech), and 10 ng/mL IL-15 (PeproTech). To evaluate functional anti-tumor activity, the T cells were harvested from culture, quantified, and co-cultured with CD19-positive Raji target cells. Raji cells were cultured in RPMI with 10% FBS (Thermo) and 1X penicillin/streptomycin (P/S) (Corning) prior to co-culture across a range of effector-to-target (E:T) ratios. Co-cultures were maintained in X-VIVO 15 medium with 5% human AB serum (MilliporeSigma), 2 mM L-glutamine, 55 μM 2-mercaptoethanol, 1X NEAA at 37 °C under 5% CO_2_ for 48 hours. Following the incubation period, cells were harvested and stained with anti-human CD19 antibodies (BioLegend) to identify surviving Raji cells. Absolute quantification of viable CD19⁺ target cells was performed using counting beads (Invitrogen) in accordance with the manufacturer’s protocol. Target cell cytotoxicity was calculated as the percentage reduction of viable CD19⁺ Raji cells relative to baseline control cultures and normalized to counting beads. Data were acquired via flow cytometry and expressed as mean ± SEM from independent biological replicates.

### LgBiT/HiBiT Intracellular Protein Detection assay

MiaPaCa-2 HiBiT-KRAS(G12C)-KI cells (Promega) were maintained in Dulbecco’s Modified Eagle Medium (DMEM) + GlutaMAX (Gibco), then plated in 6-well plates in DMEM + GlutaMAX (Gibco) supplemented with 10% FBS and 1% P/S and treated with KRAS-targeting PROTAC LC-2 (Selleck Chem) for 24 hours at 37 °C under 5% CO_2_. Cells were subsequently dissociated using 0.25% Trypsin/EDTA (Gibco) to generate a single cell suspension and boosted to deliver 3 kDa Cascade blue dextran (Thermo) and LgBiT protein (Promega) in Opti-MEM (Gibco). Delivery of dextran into live cells was quantified via flow cytometry (Attune NxT, as described above). Cells were washed with CO_2_-independent media (Gibco) before seeding in NanoGlo Endurazine (Promega) in CO_2_-independent media containing LC-2 per the manufacturer’s instructions. Cells were incubated for 2 hours at 37°C under 5% CO_2_ prior to luminescence detection on the GloMax plate reader (Promega).

### NanoBRET Target Engagement assay

HeLa cells were transfected with a Src-NanoLuciferase fusion vector (Promega) using Lipofectamine 3000 (Thermo) in Opti-MEM (Gibco) following the manufacturer’s instructions and plated in 6-well plates. After 20 hours, transfected cells were treated with cell-permeable Src kinase inhibitors (ponatinib or bosutinib; SelleckChem) for 2 hours at 37 °C under 5% CO_2_. Cells were then dissociated into a single cell suspension using 0.25% Trypsin/EDTA (Gibco), then passed through a 40 µm mesh strainer (VWR). Cells were resuspended in Opti-MEM and boosted to deliver 3 kDa dextran and a cell impermeable kinase tracer (Kinase Tracer-01, Promega) which was specifically selected to serve as an example of an impermeable assay reagent for intracellular use. Intracellular delivery of dextran to live cells was measured via flow cytometry (Attune NxT, as described above). To detect kinase tracer competition, cells were seeded in 100 µL of Opti-MEM in a white-walled 96-well plate and incubated for 2 hours at 37°C under 5% CO_2_ before adding NanoBRET Nano-Glo substrate (Promega) and NanoLuc extracellular inhibitor (Promega) per the manufacturer’s recommendations. Bioluminescence Resonance Energy Transfer (BRET) was measured on the GloMax plate reader (Promega). BRET ratio was calculated as the ratio of acceptor-to-donor luminescence, and corrected by subtracting the baseline BRET ratio of a control sample to which no kinase tracer was added.

### Intracellular Lumit immunoassay

Live-cell detection of intracellular phospho-BTK was performed using the Lumit pBTK (Tyr223) immunoassay (CS3397A20 from Promega) adapted for intact cells^19^, in which the four Lumit assay antibodies (two primaries and two NanoBiT-conjugated secondaries) were co-delivered intracellularly via mechanically-mediated poration prior to detection, in place of the conventional lysate-based workflow. For primary B cell analysis, human PBMCs were isolated from whole blood using a standard Ficoll-based protocol^20^ and B cells were stimulated within the context of the PBMC mixture with CpG oligodeoxynucleotide (InVivoGen) in ImmunoCult Human B Cell Expansion medium (STEMCELL) at a concentration of 1 x 10^6^ cells/mL at 37°C under 5% CO_2_. Following 3 days of stimulation, IL-7 (PeproTech) was added to the culture medium and cells were maintained at a concentration of 1 x 10^6^ cells/mL. On day 7, cells were harvested, counted, and re-plated a concentration of 5 x 10^6^ cells/mL in RPMI 1640 (Gibco), then subsequently rested for 3 hours at 37°C under 5% CO_2_. For Ramos cell analysis, cells were maintained in culture in RPMI (Gibco) + 10% FBS + 1% P/S at 37°C under 5% CO_2_, then harvested and re-plated as described above on the day of assay.

Following the resting period, cells were treated with 1 mM of pervanadate solution (consisting of 16 mM Na_3_VO_4_, 0.03% H_2_O_2_, in water) and incubated at 37°C under 5% CO_2_ for 40 minutes. After incubation, the cells were passed through a 40 µm mesh strainer (VWR) and boosted to deliver the four Lumit antibodies: mouse anti-BTK primary antibody, rabbit anti-phospho-BTK (Y223) primary antibody, and two Lumit secondary antibodies Lumit Anti-mouse Ab-LgBiT and Lumit Anti-Rabbit Ab-SmBiT (Promega). For Lumit detection, cells were plated in a white-walled 96-well plate and incubated for 20 minutes at RT. After incubation, Lumit Detection Reagent, consisting of Furimazine substrate diluted in Immunoassay Reaction Buffer (Promega) was added to the cells following the manufacturer’s instructions. The plate was agitated at 400 rpm for 2 minutes and read using the preset Lumit Immunoassay settings on the Promega GloMax plate reader. Raw luminescence values were recorded and compared between treated and untreated cell groups. Baseline control samples were prepared using cell lysate following the manufacturer’s instructions for a conventional Lumit Immunoassay without processing cells through the micro-pore array.

### Multi-modal T cell engineering at scale

To evaluate mechanically-mediated poration within a cell-therapy-type workflow, unstimulated human T cells were isolated from human PBMCs as described above and boosted at research scale to deliver β2-microglobulin (B2M) CRISPR RNP complexes (IDT). Cell concentration and viability were determined using a NC-202 (ChemoMetec) with AO/PI cartridge (ChemiMetec). Viable cell counts were used to adjust the cell suspension to 1.25 x 10^6^ viable cells/mL before seeding into G-Rex 6M plates (ScaleReady) in CTS OpTmizer serum-free medium (Thermo) supplemented with CTS OpTmizer expansion supplement (Thermo), 5% human AB serum (MilliporeSigma), 2 mM L-glutamine, 1% P/S, 15 ng/mL IL-7 (Peprotech), and 5 ng/mL IL-15 (Peprotech). T cells were activated with TransAct CD3/CD28 reagent (Miltenyi Biotec) and expanded in G-Rex plates for 9 days at 37°C under 5% CO_2_ following the manufacturer’s recommendations. Activated T cells were subsequently boosted to deliver *GFP* mRNA (TriLink) at clinical scale using the LOVO cell processing system (Fresenius Kabi) as the driving force. Following this delivery, cells were cultured in the above media for 48 hours before assessing GFP expression and B2M protein knockout efficiency by flow cytometry as described.

### PBMC and T cell engineering at scale

Human PBMCs or unstimulated T cells were isolated from whole blood as described above, mixed with cargo and boosted to deliver dextran and B2M CRISPR RNPs as described above, using the CTS Rotea Counterflow Centrifugation System (Thermo Fisher Scientific) as the driving force. Cells were collected and immediately evaluated for dextran delivery via flow cytometry as described, or cultured prior to flow cytometry analysis for 48 hours at 37 °C under 5% CO_2_ in X-VIVO 15 medium with 5% human AB serum (MilliporeSigma), 2 mM L-glutamine, 55 μM 2-mercaptoethanol, 1X NEAA, 200 U/mL IL-2, 10 ng/mL IL-7 (PeproTech), and 10 ng/mL IL-15 (PeproTech).

### T cell engineering in whole blood

Peripheral whole blood was collected from healthy donors and processed under sterile conditions. CircRNAs encoding mbIL-2 and a CD19-CAR were boosted directly into whole blood using a pore geometry optimized for unstimulated human T cells. Following poration, cells were seeded into 12-well culture plates and cultured at 37 °C under 5% CO₂ in X-VIVO 15 medium (Lonza) supplemented with 5% human AB serum (MilliporeSigma), 2 mM L-glutamine, 1X NEAA, and 55 μM 2-mercaptoethanol. Expression of mbIL-2 and CD19 CAR were assessed by flow cytometry as described 40 hours following mechanically-mediated poration.

## Acknowledgements

The authors wish to thank Michael Finot, Pedro Duarte, Cole Constantineau, Collin Mason, and Gabriel Kornilowicz for their valuable input on system design and prototyping. We thank Dr. Orion Weiner and Nitya Kopparapu at UC San Francisco for their contributions toward the neutrophil migration assay and the Smith lab at University of Illinois Urbana-Champaign for their contributions toward quantum dot delivery to PBMCs. We thank Matt Robers and the team at Promega for assistance, consultation, and sharing of reagents and protocols for target engagement assays. We thank the teams at ScaleReady, Fresenius Kabi, and Thermo Fisher Scientific for their assistance in the design and implementation of the T cell engineering clinical applications, including instrument training, and sharing of equipment and materials required to complete the work presented.

---

**Extended Data Figure 1.**
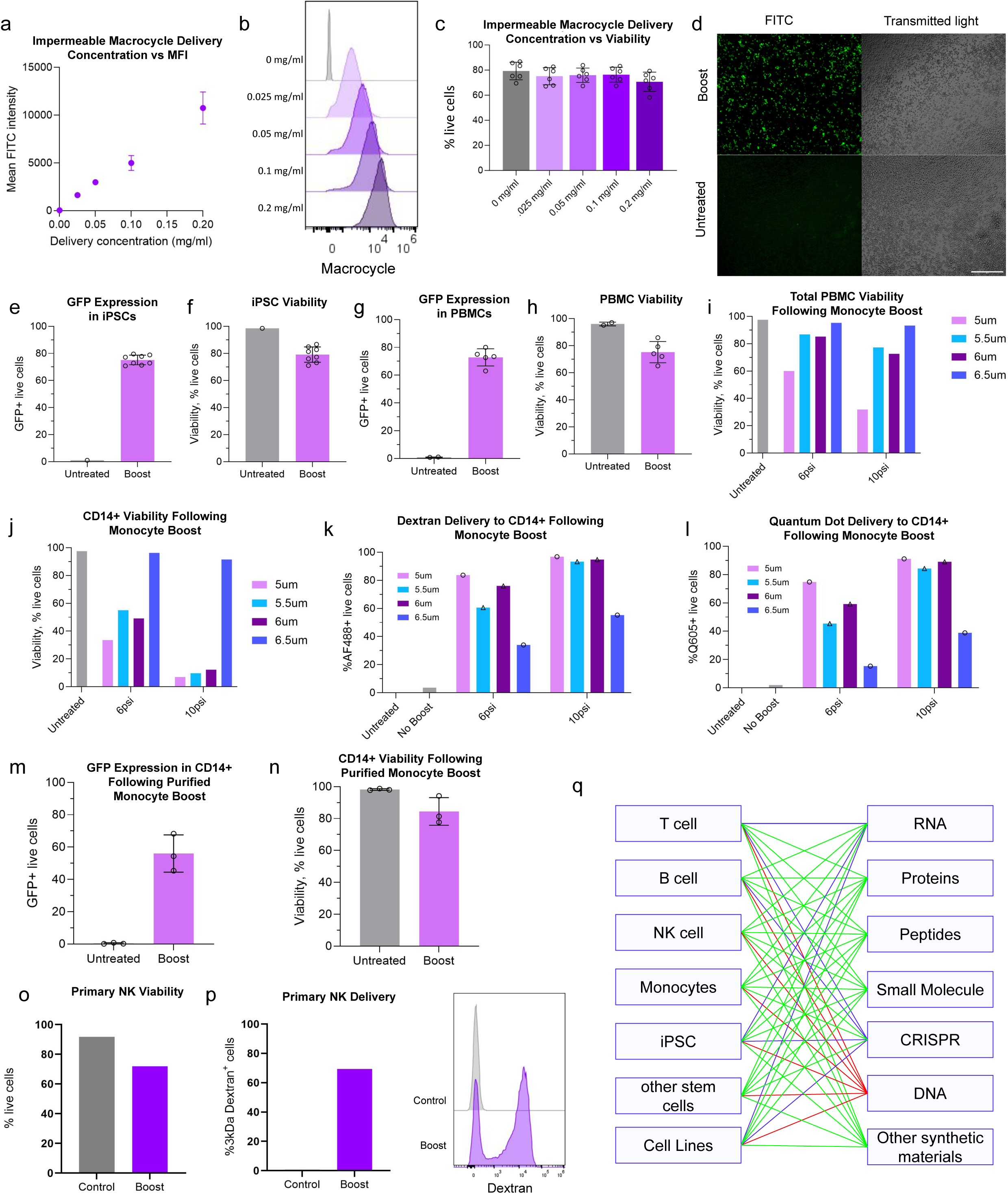
Consistent performance of mechanically-mediated poration across diverse cell lineages. **a-c**, Linear and stoichiometric correlation of impermeable macrocycle delivery to HeLa cells, demonstrating increasing fluorescence intensity scaling with increasing cargo concentration (**a**,**b**) and post-poration viability (**c**) (n = 6 from two independent experiments). **d**, Microscopy depicting diffuse distribution of macrocycle in HeLa cells. Scale bar, 300 μm. **e-p**, Benchmarking of delivery efficiencies and post-poration viabilities across representative cell lines, stem cells, and primary immune cells. **e**,**f**, Representative GFP expression (**e**) and viability (**f**) in human iPSCs following *GFP* mRNA delivery (n = 8) comparing untreated control cells against platform-porated (“boost”) conditions. **g**,**h**, Representative GFP expression (**g**) and viability (**h**) in human PBMCs following *GFP* mRNA delivery (n = 5 biological replicates). **i**-**l** Optimization matrix for targeted monocyte delivery within mixed PBMC populations, showing total PBMC viability (**i**), monocyte viability (**j**), 3 kDa dextran delivery to CD14+ population (**k**) and quantum dot intracellular accumulation in CD14+ (**l**). **m**,**n**, GFP expression (**m**) and post-poration viability (**n**) in purified CD14+ monocytes following *GFP* mRNA delivery. **o**,**p**, Post-delivery viability (**o**) and 3 kDa dextran delivery (**p**) in primary human NK cells. **q**, Comparative performance matrix of mechanical delivery against conventional biological and physical delivery modalities. Green denotes a clear functional advantage over traditional methods, including historically intractable macromolecules; purple indicates comparable performance thresholds to established methods; red highlights applications showing no significant performance differentiation.

**Extended Data Figure 2.**
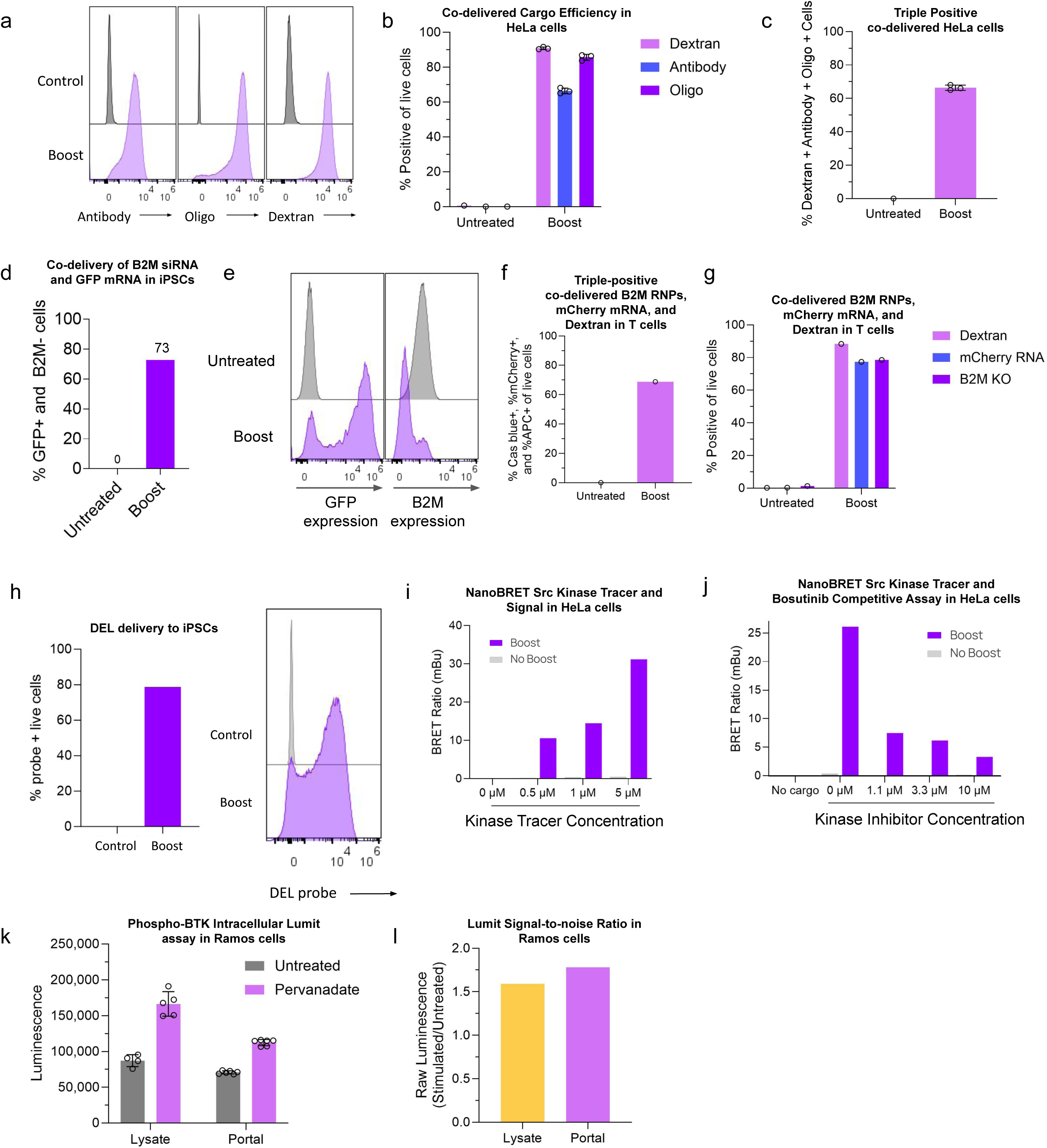
Robust cell engineering opportunities across cell lineages and cargo modalities. **a**-**c**, Simultaneous co-delivery of an antibody, oligonucleotide, and dextran in HeLa cells demonstrating single-step, multi-modal cell engineering with preserved cargo stoichiometry. **a**, Representative flow cytometry histograms of intracellular antibody, oligonucleotide, and dextran following boost. **b**, Independent delivery efficiency quantification for each distinct cargo modality. **c**, Triple-positive engineering efficiency tracks with the efficiency of the least-efficient cargo. **d**,**e**, Co-delivery of *B2M*-targeted siRNA and *GFP* mRNA to human iPSCs via the platform yielding highly efficient multi-modal modulation (**d**). **e**, Representative flow cytometry histograms displaying resultant GFP expression and residual surface B2M levels. **f**,**g**, Poration-mediated co-delivery of 3 kDa dextran, *mCherry* mRNA, and CRISPR RNPs targeting the *B2M* locus in unstimulated human T cells, showing triple positive multiplexing efficiency (**f**) and corresponding single-positive efficiencies for each modality (**g**). **h**, Intracellular delivery of DELs into human iPSCs via the platform. **i**,**j**, Live-cell NanoBRET target engagement kinetics in HeLa cells. Data show a dose-dependent BRET signal increase scaling with the initial platform-loaded concentration of a traditionally impermeable Src kinase tracer (**i**), followed by a competitive, dose-dependent decrease in signal upon exposure to Src kinase inhibitor Bosutinib (**j**). **k**,**l**, Live-cell intracellular phospho-BTK detection in Ramos cells. Mechanical delivery of four phospho-BTK antibodies following pervanadate stimulation yields a robust intracellular phosphorylation signal (**k**) with a signal-to-noise ratio comparable to traditional lysate-based detection (**l**).

**Extended Data Figure 3.**
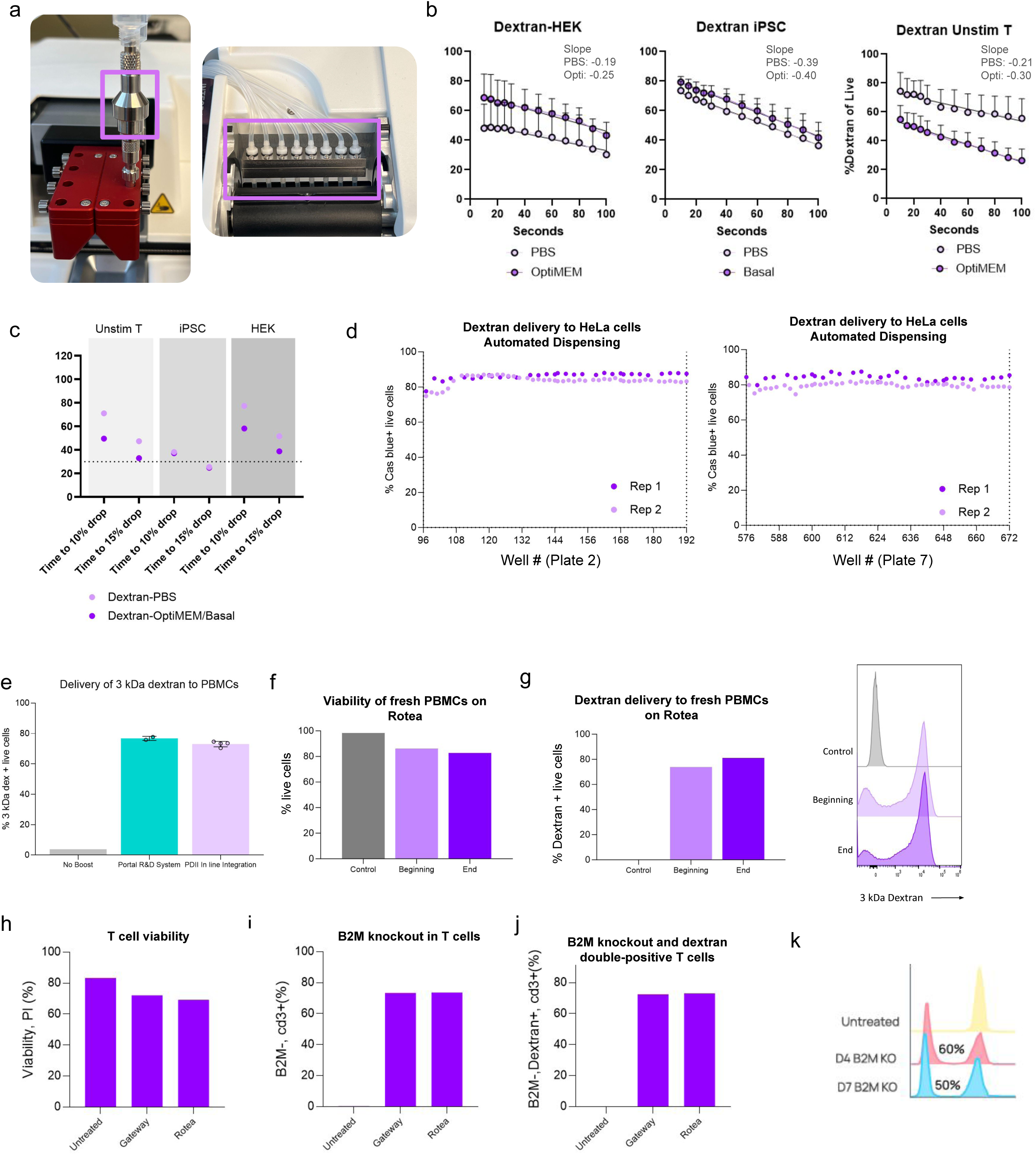
Fluidic automation compatibility and post-poration membrane closure kinetics. **a**, Integration of the mechanically-mediated poration platform with automated multichannel and peristaltic pump dispensing instrumentation, illustrating representative hardware configurations utilizing the Nnano/Gyger Certus Flex and Integra WellJet systems. **b**,**c**, Membrane closure kinetics evaluated via time-delay 3 kDa dextran delivery to HEK, iPSC, and human T cells following boost. Data demonstrate a time-dependent decay in cytosolic uptake (**b**), where delivery efficiency is maintained within 15% of the baseline simultaneous (pre-mixed) delivery efficiency (time = 0, cargo delivered during pore transit) within a 30 second post-poration window (**c**). **d**, High throughput (HT) delivery efficiency across plates 2 and 7 in a continuous automated dispensing of HeLa cells with platform integration on the GNF One-Tip Dispenser into plates pre-loaded with 3 kDa dextran. **e**, Comparative efficiencies of 3 kDa dextran delivered to human PBMCs via mechanically-mediated poration, contrasting conventional pre-mixed cargo on the research-scale architecture against post-poration loading via platform-integrated PDII automated dispensing into a pre-loaded multiwell plate. **f**,**g**, Scaled preliminary workflow delivering dextran to fresh human PBMCs using an integrated cartridge on Thermo’s Rotea cell processing instrument. **f**, Viability was consistently maintained from beginning to end of the processing run, with high delivery efficiency to all cells throughout the run from beginning to end (**g**). **h**-**j**, Scaled preliminary editing in primary human T cells on the Rotea demonstrating equivalent viability (**h**), B2M KO (**i**), and double-positive edited, dextran-labeled cells (**j**) as compared to delivery at research scale on Portal’s Gateway instrument. **k**, Longitudinal *B2M* KO efficiency measured during primary human T cell expansion following mechanically-mediated poration at research-scale in the workflow outlined in Fig. 3m.

**Supplementary Video 1.**
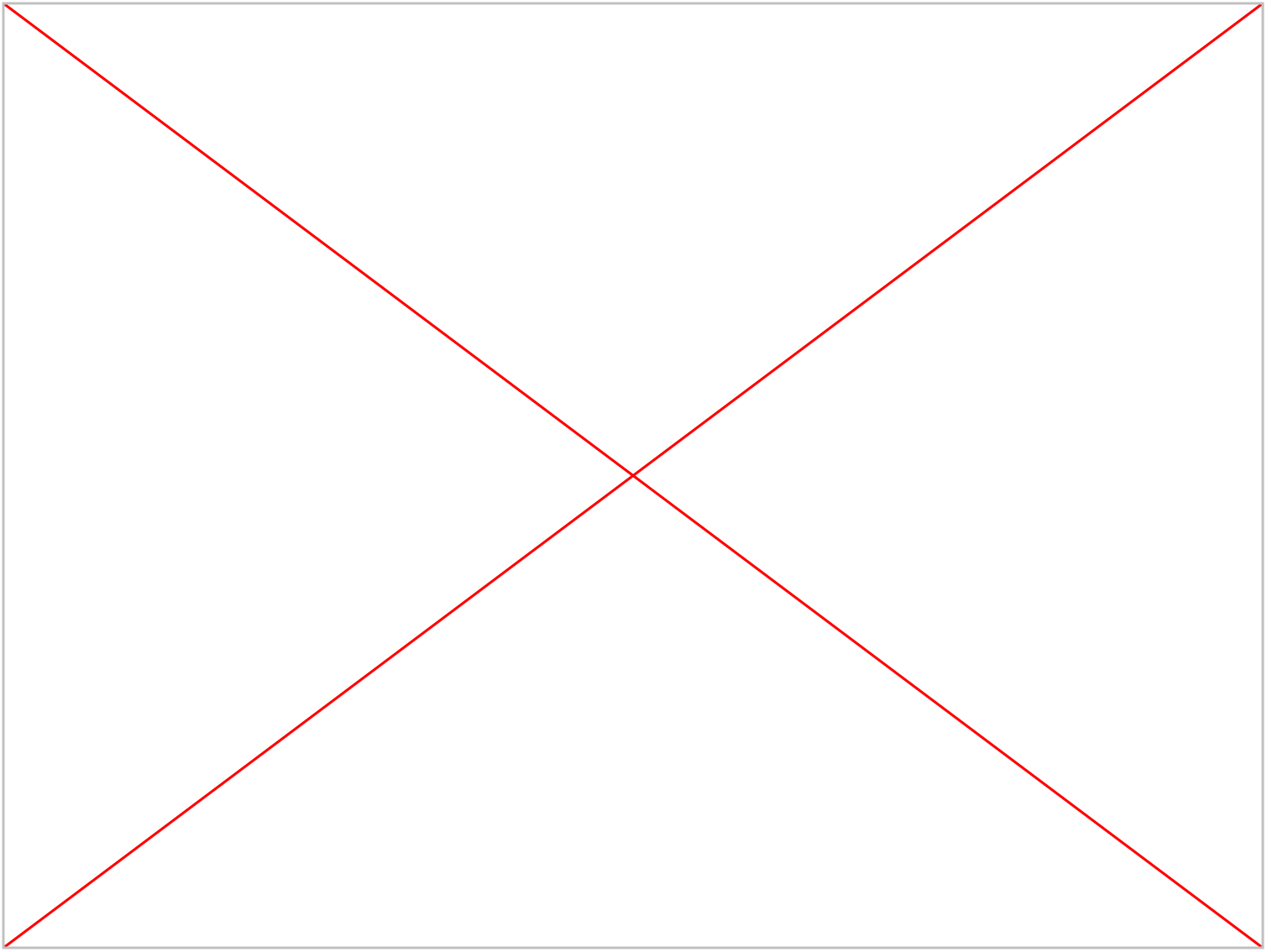
Migration dynamics of primary human neutrophils following *GFP* mRNA delivery. Time-lapse fluorescence microscopy of primary human neutrophils following mechanically-mediated delivery of *GFP* circRNA. Cells exhibit robust GFP expression, homeostatic morphology, and migration on a fibronectin-coated plate in the presence of chemokine introduced into the media. Images were captured every 5 s.

**Supplementary Video 2.**
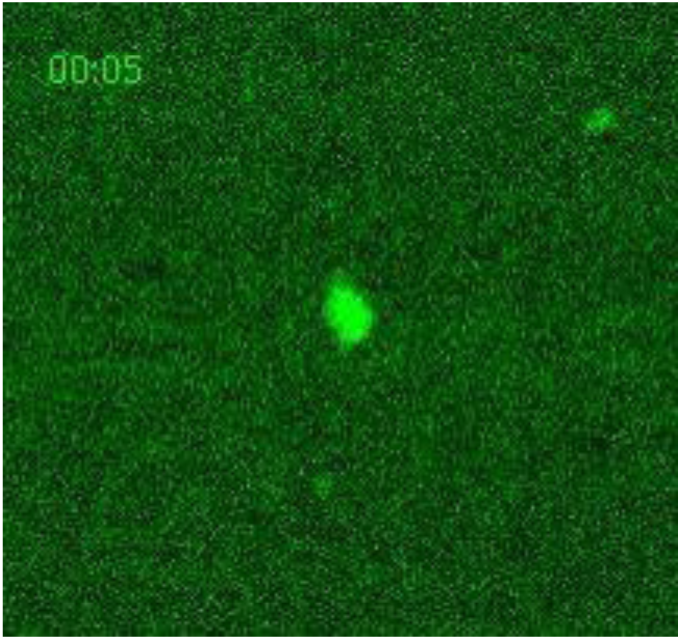
High-magnification tracking of an engineered human neutrophil. High-magnification time-lapse sequence tracking a single GFP-expressing primary human neutrophil actively migrating in response to a chemokine present in the media following mechanically-mediated *GFP* circRNA delivery. Images were captured every 5 s.

